# Maternal immune activation alters social affective behavior and sensitivity to corticotropin releasing factor in male but not female rats

**DOI:** 10.1101/2022.07.05.498833

**Authors:** Nathaniel S. Rieger, Alexandra J. Ng, Shanon Lee, Bridget H. Brady, John P. Christianson

## Abstract

Prenatal infection increases risk for neurodevelopmental disorders such as autism in offspring. In the rodents, prenatal administration of the viral mimic Polyinosinic:polycytidylic acid (Poly I:C) allows for investigation of developmental consequences of gestational sickness on offspring social behavior and neural circuit function. Because maternal immune activation (MIA) disrupts cortical development and sociability, we examined social decision-making in a rat social affective preference (SAP) task. Following Poly I:C (0.5 mg/kg) on gestational day 12.5, male adult offspring (PN 50) exhibited atypical social interactions with stressed conspecifics whereas female SAP behavior was unaffected by maternal Poly I:C. Social responses to stressed conspecifics depend upon the insular cortex where corticotropin releasing factor (CRF) modulates synaptic transmission and SAP behavior. We characterized insular field excitatory postsynaptic potentials (fEPSP) in adult offspring of MIA or control treated dams. Male MIA offspring showed decreased sensitivity to CRF (300 nM) while female MIA offspring showed greater sensitivity to CRF compared to sham offspring. These sex specific effects appear to be behaviorally relevant as CRF injected into the insula of male and female rats prior to social exploration testing had no effect in MIA male offspring but increased social interaction in female MIA offspring. We examined the cellular distribution of CRF receptor mRNA but found no effect of maternal Poly I:C in the insula. Together these experiments reveal sex specific effects of prenatal infection on offspring social decision making and identify insular CRF signaling as a novel neurobiological substrate for autism risk.

## Introduction

Epidemiological studies reveal significant increases in the prevalence of neuropsychiatric disorders such as autism spectrum disorders (ASD) and schizophrenia following viral pandemic outbreaks (Estes and McAllister, 2016) and prenatal inflammation is thought to play a role in risk (Careaga et al., 2017). Maternal infections during gestation increase cytokine levels in offspring, activating toll-like receptors and leading to anatomical changes, such as cortical thinning, and behavioral changes, such as decreased sociability and communication deficits, indicative of disorders like ASD (Bilbo et al., 2018; Careaga et al., 2017; Garbett et al., 2012; Lombardo et al., 2018; Malkova et al., 2012).These infections can be modeled in rodents in a maternal immune activation model (MIA) (Estes and McAllister, 2016). MIA is achieved through the induction of fever and inflammation over a ∼48h period in gestating mothers via i.p. injection of endotoxins or viral mimics (polyinosinic: polycytidylic acid; Poly I:C) (Hsiao and Patterson, 2011; Kentner et al., 2019) that increase inflammatory cytokines and chemokines (Williamson et al., 2011). Interestingly, the Poly I:C model of MIA reveals sex differences in behavioral outcomes, including sociality (Bilbo et al., 2018; Carlezon et al., 2019; Heuer et al., 2019; Howerton and Bale, 2012; Hsiao and Patterson, 2011). In general, behavioral outcomes and neural alterations of MIA in rodents are more pronounced in males compared to females (Haida et al., 2019). However, the underlying neurobiology of why males are more likely to develop behavior deficits but females are buffered from these effects is not well understood.

Mechanistic studies of MIA in mice identified anatomical and functional changes in subcortical regions (Bilbo et al., 2018; Heuer et al., 2019; Howerton and Bale, 2012). Behaviorally, MIA reduces approach to social novelty and affiliative behavior which seems to reflect alterations in the way social stimuli engage the mesolimbic reward system (Aguilar-Valles et al., 2020; Straley et al., 2017; Vitor-Vieira et al., 2021). We sought to investigate social behaviors that are context dependent, specifically whether to approach or avoid conspecifics that received a stressor. The choice to approach or avoid a conspecific in distress requires cortical structures like the anterior cingulate and insular cortex which regulate social approach, in part, via interaction with the mesolimbic reward system (Rogers-Carter et al., 2019; Rogers-Carter and Christianson, 2019). The insula is a site of interest as a link between MIA and social dysfunction because in both ASD and schizophrenia, abnormal insula activity and functional connectivity are frequently reported correlates of social cognition (Baribeau and Anagnostou, 2015; Hogeveen et al., 2018; Odriozola et al., 2016). Further, the insula appears to encode some aspects of immune activation (Koren et al., 2021). The insula’s role in social cognition emerges from its importance to processing interoceptive and external stimuli (Esan et al., 2015) allowing for the evaluation of social and emotional cues. The insula receives a number of sensory afferents (Augustine, 1996) and is reciprocally connected to several cortical and subcortical regions (Nieuwenhuys, 2012). In this way, the insula is said to be part of a ‘salience network’ (Gogolla, 2017; Uddin and Menon, 2009) which is vital to integrating external and internal cues and the insula is hypothesized to be a hub among larger networks necessary for controlling social decision-making and emotional processing (Ebisch et al., 2011). Accordingly, we hypothesized that insula function, and insula-dependent social behaviors, would be vulnerable to change as a result of MIA.

To begin our investigation of MIA effects on insular function we focused on corticotropin releasing factor (CRF). CRF is an organizational peptide principly known for its role in initiating the stress response (Smelik, 1960), but it is also involved with coordinating social behavior (Hostetler and Ryabinin, 2013). CRF plays a role in evaluating external negative cues and adjusting social responses to match these cues (Hostetler and Ryabinin, 2013; Hupalo et al., 2019) and CRF in the periventricular nucleus of the hypothalamus (PVN) is necessary for the transmission of emotional states between rodents (Sterley and Bains, 2021; Sterley et al., 2018). Behavior in SAP tests, for male rats, requires CRF which shapes insular activity by augmenting excitatory synaptic transmission in a CRF receptor 1 (CRF_1_) dependent fashion (Rieger et al., 2022). CRF signaling is drastically altered by early life stress in subcortical areas (Authement et al., 2018) and it has been suggested that MIA might influence neural-immune and CRF systems (Bale et al., 2010; Bilbo and Schwarz, 2012) leading us to speculate that MIA could also influence insular CRF function.

This project investigated how maternal immune activation influenced social affective behavior, insular cortex (IC) and the stress-related neuropeptide corticotropin-releasing factor (CRF) which convening lines of research (Arakawa et al., 2011; Chen and Hong, 2018; Felix-Ortiz and Tye, 2014; Hatfield et al., 1993; Kiyokawa, 2015; Parkinson and Simons, 2009; Rieger et al., 2022; Sterley et al., 2018) implicate in the influence of social stress cues on observer behavior. We hypothesized that MIA would alter insular CRF function and cause aberrations in social decision making. Dams received Poly I:C to generate MIA male and female offspring and we compared their social decision making to saline treated controls using the SAP test. In separate experiments, we tested for changes in insular CRF sensitivity in slice physiology using a multiple electrode array and in behavior using one-on-one social interaction tests. Finally, we tested for changes in insular CRF_1_ mRNA distribution using fluorescent *in situ* hybridization.

## Methods

### Animals and Breeding

Adult male and female rats (PN 45) were obtained from Charles River Laboratories (Wilmington, MA) and acclimated to the vivarium for 2 weeks prior to breeding. Rats were housed in standard cages of 2 same-sex conspecifics and given access to food and water *ad libitum*. The vivarium was maintained on a 12 h light dark cycle. Breeding occurred overnight during the dark cycle and all behavioral testing took place within the first 4 h of the light cycle. Breeding followed standard procedures (Kentner et al., 2019). One male and one female were housed together overnight in a standard cage for up to 5 nights. Each morning females were checked for the presence of a sperm plug. The presence of a sperm plug was used as the start of the gestation timeline (E 0.5). Females were weighed daily throughout pregnancy and Poly I:C or saline injections occurred on E 12.5 (Han et al., 2011). All procedures and animal care were conducted in accordance with the NIH *Guide for the care and use of laboratory animals* and all experimental procedures were approved by the Boston College Institutional Animal Care and Use Committee.

### Maternal Immune Activation

Poly I:C (Sigma-Aldrich, St. Louis, MO, Product #P9582, Lot #118M4035V) was dissolved in saline to a dose of 0.5 mg/kg. This dose was selected as the minimum dose necessary to induce a cytokine and fever response in dams (Fortier et al., 2004) and a behavioral effect in adult offspring (Han et al., 2011). Poly I:C can be contaminated with endotoxin and we did not test for this, as such the behavioral changes we see could be due to bacterial infection as well as the by the viral action of Poly I:C. Poly I:C or saline was injected i.p. to pregnant dams on day E 12.5. Rectal temperature of dams was taken prior to injection as a baseline and again at 4, 8 and 24 hours post injection to monitor for fevers. Over the course of 48 h post injection dams were also monitored for behavioral signs of sickness including piloerection, drooping eyelids and malaise. Pregnancies were then allowed to continue to parturition. Post birth, dams were allowed to raise their pups normally and were monitored for maternal care including huddling and grooming once a week until weaning (PN 21) with no differences being detected. Offspring were then randomly assigned to behavioral tests (max 2 per litter) or electrophysiological studies (max 1 per litter) and testing occurred on PN 60. Statistics were run at the level of the individual.

### Social Affective Preference Test (SAP)

In the SAP test, rats exhibit preferences in social interaction with conspecifics depending upon the age and socioemotional condition of the targets. SAP tests were performed exactly as previously reported (Rieger et al., 2022; Rogers-Carter et al., 2018). The test involves observing the social interaction behavior of a focal test rat provided with 2 unfamiliar conspecifics, one of which underwent a mild footshock stress prior to the test and the other naive to treatment. Male and female rats typically avoid stressed adult conspecifics but will approach juvenile stressed conspecifics (Rogers-Carter et al., 2018). SAP testing occurred over 3 days with days 1 and 2 serving as habituation days and day 3 serving as the test day (Fig 4). On day 1 rats were placed into the testing chamber (50 cm x 40 cm x 20 cm) for 1h. On day 2, rats were placed into the chamber for 1h with 2 naive juvenile or adult same-sex conspecifics. Conspecifics were placed at opposite ends of the testing chamber in a 18 cm x 21 cm x 10 cm cage with horizontal bars placed 1 cm apart on one side to facilitate interactions. Test rats were then allowed to freely interact for 5 min. Time spent investigating via nose-nose and nose-body sniffing and reaching for the conspecific was quantified. On day 3 rats were placed in the chamber 1hr before testing with a naive and stressed conspecific. Stressed conspecifics were exposed to 2, 5s 1 mA footshocks (Colburn Instruments) prior to testing. Quantification was done as above and scored by an observer blind to treatment. Tests were also video recorded and scored by a second observer to determine interrater reliability.

### Electrophysiology

#### Solutions

Standard solutions were used for artificial cerebrospinal fluid (aCSF) as previously (Rieger et al., 2022; Rogers-Carter et al., 2018) and all reagents were purchased from Fisher, Sigma or Tocris. aCSF recording composition was (in mM) NaCl 125, KCl 2.5, NaHCO_3_ 25, NaH_2_PO_4_ 1.25, MgCl_2_ 1, CaCl_2_ 2 and Glucose 10; pH = 7.40; 310 mOsm; aCSF cutting solution was: Sucrose 75, NaCl 87, KCl 2.5, NaHCO_3_ 25, NaH_2_PO_4_ 1.25, MgCl_2_ 7, CaCl_2_ 0.5, Glucose 25 and Kynureinic acid 1; pH=7.40, 312 mOsm.

### Insular cortex slices

Adult male and female rats were anesthetized by isoflurane, intracardially perfused with chilled (4°C) cutting solution and rapidly decapitated. The brain was sliced on a vibratome (VT-1000S, Leica Microsystems, Nussloch, Germany) in 300 μm coronal segments containing the insula. Slices were then transferred to oxygenated (95% O_2_, 5% CO_2_) aCSF at 37°C for 30 minutes followed by 30 minutes at room temperature prior to any electrophysiological recordings.

### Evoked Field Excitatory Postsynaptic Potentials

Evoked field excitatory postsynaptic potentials (fEPSPs) were recorded on a 6 × 10 perforated multielectrode array (MCSMEA-S4-GR with 60pMEA100/30iR-Ti Array, Multichannel Systems) with integrated acquisition hardware (MCSUSB60) and analyzed with MC_RACK software (version 3.9) exactly as described previously (Rieger et al., 2022; Rogers-Carter et al., 2018). Slices were placed onto the array and affixed by downward suction through the perforated array. CRF (300 nM), the dose found to previously augment fEPSPs (Reiger et al., 2022), was dissolved in water, diluted to its final concentration in aCSF and bath applied to slices at 37 °C.

A stimulating electrode was selected in the insular cortex and fEPSPs I/O curves were recorded in the adjacent electrodes following electrical stimulation before (aCSF), and 10 min after CRF application. Stimulations ranged from 0-5 V and occurred in biphasic (220 μs) 500 mV increments in triplicate. fEPSPs that showed clear synaptic responses and occurred close to the stimulating electrode were normalized to the maximum 5 V response of the slice at baseline and channels from the same slice were averaged for group analysis across subjects. Slices received only one drug treatment.

### Surgical implantation of indwelling cannula

While under isoflurane anesthesia (2-5% v/v in O_2_), rats were surgically implanted with bilateral guide cannula (26-gauge, Plastics One, Roanoke VA) in the insular cortex (from bregma: AP: -1.8 mm, M/L: ± 6.5 mm, D/V: -6.8 mm from skull surface) that were affixed with stainless steel screws and acrylic dental cement. Immediately following surgery, rats were injected with analgesic meloxicam (1 mg/kg, Eloxiject, Henry Schein), antibiotic penicillin (12,000 units, Combi-pen 48, Covetrus) and Ringer’s solution (10 mL, Covetrus). Rats were then returned to their homecage and allowed 7-14 days for recovery prior to behavioral testing.

CRF (Tocris) was first dissolved in DI water and diluted to 300 nM concentration in a vehicle of 0.9% saline. Injections were 0.5 μL/side for all drugs and were infused at a rate of 1.0 μL/minute with an additional one minute diffusion time after injection. After behavioral testing concluded, rats were overdosed with tribromoethanol (Sigma) and brains were immediately dissected, flash frozen on dry ice, and sectioned on a cryostat at 40 μm. Slices were then mounted onto gelatin-subbed slides (Fisher) and stained with cresyl-violet to verify cannula placements by comparing microinjector tip location to the rat whole brain stereotaxic atlas. Rats were excluded from all analyses if their cannula placements were found to be outside the insula or if cannulas were occluded prior to injection.

### Social Exploration

One-on-one social exploration tests were conducted to determine differences in overall social behavior and whether this was altered based on MIA and insular CRF. Test animals were placed in a standard cage (18 cm x 24 cm x 18 cm) for 1h. 40 min prior to testing animals were microinjected with 300 nM CRF in the insula, a dose shown to increase social interaction in male rats (Rieger et al., 2022). At the start of testing, a juvenile same-sex conspecific was placed in the chamber with the test rat and they were allowed to freely interact for five minutes. Time spent investigating via sniffing and grabbing of the conspecific was quantified by an observer blind to treatment and video recorded. Rats were tested in a within-subject design on 2 consecutive days such that rats randomly received either vehicle or CRF on day one and the opposite on day two; treatment order was counterbalanced. Conspecifics were used twice per day, but no test rat interacted with the same conspecific twice.

### RNAscope *in situ* Fluorescent hybridization

Coronal sections containing the posterior insular cortex (Bregma -1.8) were collected on a freezing cryostat at 20μm thick and mounted to SuperFrost+ slides and stored at -80C until processing. RNAScope was performed according to the vendor’s instructions (ACDBio). Briefly, tissue was thawed, fixed and treated with a RNAScope cocktail including probes for CRF1 (*crhr1*, catalog #318911), GAD (*gad1*, catalog #), vesicular glutamate transporter 1 (*vglut1, catalog #317001*) and DAPI. Probes were amplified and visualized with AMP1, AMP2, AMP3, AMP4, and DAPI, coverslipped with aqueous antifade media (Prolong Gold) and imaged on a Zeiss AxioImager Z2 microscope with a digital CCD camera (ORCA 3, Hamamatsu) using an Apotome2 and 20x objective (N.A. = 0.8) and fluorescent filter cubes for DAPI (365 nm excitation, Zeiss filter 49), GFP (470/40 nm excitation, Zeiss filter 38 HE), DSRed (545/25 nm excitation, Zeiss filter 43 HE) and Cy5 (640/30 nm excitation, Zeiss filter 50). All image acquisition parameters (exposure, camera gain, and display curves) were consistent for all samples. A series of multiplex, tiled mosaic images consisting of 9 z-series images per channel were stitched, deconvolved and maximum projections were saved for analysis in ImageJ. All channels were converted to binary and DAPI cells and vglut1, crhr1 and gad grains were detected using the particle counter tool. Trained observers counted the number of DAPI nuclei that were colocalized (a minimum of 3 overlapping, or adjacent grains) with each of the targets. The total number of nuclei (DAPI), glutamate cells (Vglut+DAPI), GABA cells (GAD+DAPI) and cells colocalized with crhr1, were determined). Cell counts were normalized by the total number of DAPI, DAPI+vglut, or DAPI+gad cells to compare the distribution of mRNAs with 2 way ANOVAs with sex as a between groups factor and side as a within-subject factor. The mean of left and right hemisphere counts are shown.

### Statistical Analysis

All statistics were computed in Prism (Graphpad, Version 9.2). Sample sizes were determined based on previous experiments and *a priori* power analysis. No more than 2 rats per litter were used in any given test. Subjects were randomly assigned to experimental tests and dams were randomly assigned to receive Saline or Poly I:C. All experiments utilized a counterbalanced design. To compare differences between mean scores of social interaction and electrophysiological endpoints we used analysis of variance (ANOVA). Individual replicate data are provided in the figures. Data were checked for normality and final sample sizes are indicated in the Figure Legends. In most experiments, there were within-subjects variables, which were treated as such in the analysis (repeated measures ANOVA). Main effects and interactions were deemed significant when p < 0.05 and all reported post hoc test *p* values were Sidak-adjusted to maintain an experiment-wise risk of type I errors at a = 0.05.

## Results

### Maternal immune activation

To induce MIA, pregnant dams were injected with 0.5 mg/kg Poly I:C or saline on embryonic day 12.5 (Fig. 1A). Rectal temperatures of all dams in the study were taken prior to injection (baseline) and 4 and 24 h following injection. Females injected with Poly I:C showed significantly increased temperatures and signs of sickness including piloerection and drooping eyes within 4 h of injection (Time x Treatment ANOVA F _2,20_ = 8.810, p =0.002, *η*^2^ = 0.18). The fever and sickness signs resolved within 24 h and dams continued their pregnancy normally (Fig. 1B). Only offspring of Poly I:C injected dams that had a febrile response were included in behavioral testing. Time to birth and litter size did not differ between saline and Poly I:C dams and maternal care also did not differ in huddling time which was assessed via scan sampling during the first week post-birth. Because changes in maternal behavior were not detected, offspring were raised by their own mother and not cross-fostered. Weight at weaning (PN21) did not differ significantly between offspring of Poly I:C or saline mothers.

**Figure 1:**
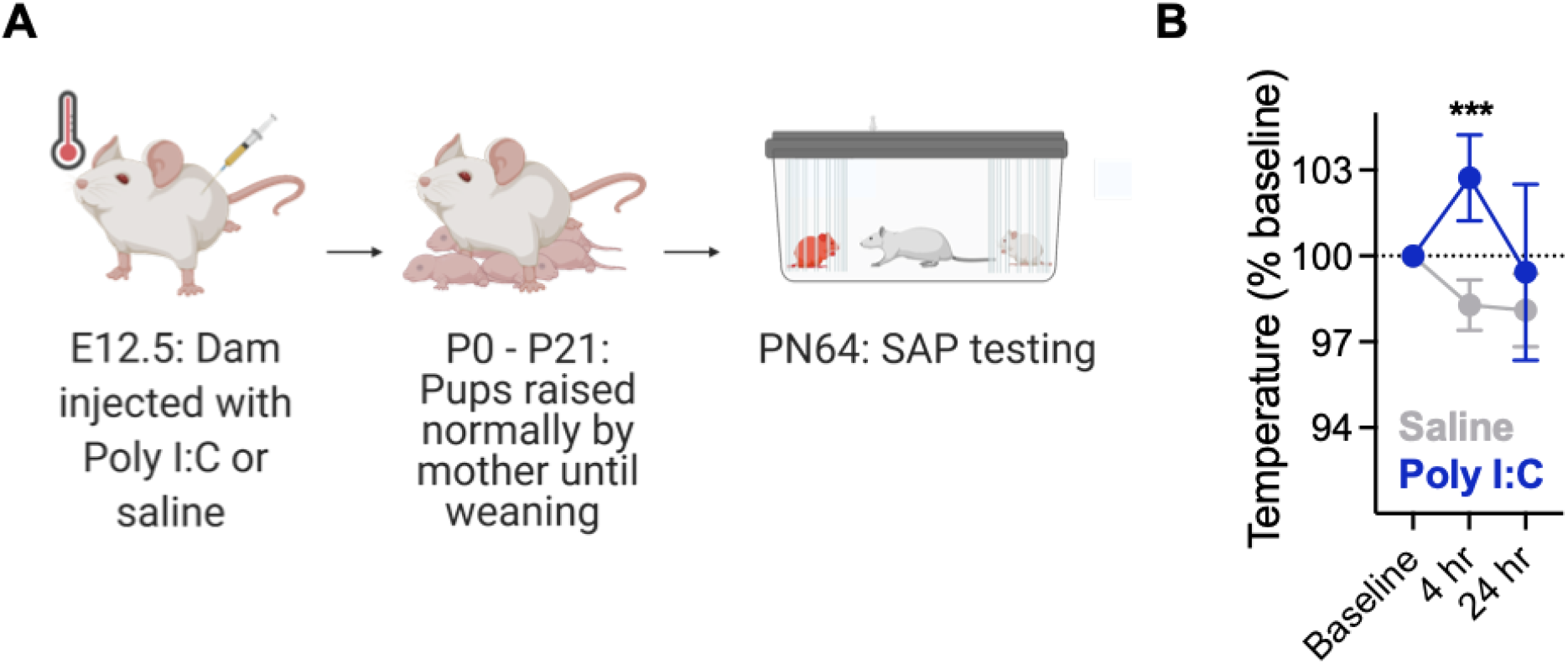
Maternal immune activation and fever verification. **A**. Timeline of MIA experiments. Dams were mated with a male partner overnight and checked for the presence of a sperm plug in the morning. Positive identification of the sperm plug was treated as day 0.5. On embryonic day 12.5 dams were injected with either 0.5 mg/kg Poly I:C or saline and allowed to return to their cages. The presence of a fever response was verified by taking temperatures at baseline (prior to injection) and 4 and 24 hours post injection. Mothers carried out their pregnancy until parturition and were allowed to raise their pups normally. Offspring of Poly I:C and saline treated mothers were randomly assigned to treatment groups and cannula implantation occured on PN50. Behavioral testing began on PN64. **B**. Mean (+/-) SEM) rectal temperatures. Mothers treated with Poly I:C developed significant fever responses with temperatures significantly increased in dams treated with Poly I:C compared to saline injected dams at 4h post injection. ***p < 0.001.

### Maternal immune activation disrupts social affective preference in males but not females

At PN60 males and females were tested for social affective preference (SAP) with juvenile (PN30) or adult (PN50) conspecifics. To assess sociability, we tallied the total time interacting on day 2 in which experimental rats were presented with 2 naive conspecifics as part of the habituation for SAP tests (Fig. 2A). Male rats, regardless of MIA treatment, spent more time investigating juvenile conspecifics (Sex by Age interaction, F(1, 76) = 10.25, P = 0.002, *η*^2^ = 0.04). MIA did not effect sociability and MIA did not result in any significant interactions with Sex or Age. On day 3 of the SAP test experimental rats were presented with one naive and one stressed conspecific. Control males, control females, and MIA females spent more time investigating stressed PN30 conspecifics (Fig. 2B, MIA by Sex by Stress interaction F(1, 75) = 14.13, p = 0.0003, *η*^2^ = 0.04, posthoc comparisons *p*s < 0.001, Sidak Corrected) and less time investigating stressed PN50 conspecifics (MIA by Sex by Age interaction F(1, 75) = 5.103, p = 0.027, *η*^2^ = 0.02, posthoc comparisons *p*s < 0.05, Sidak Corrected), male offspring of Poly I:C treated dams did not differentiate between stressed conspecifics regardless of age. To facilitate comparison across treatments, time spent interacting with the stressed conspecifics was converted to a preference score (Preference for Stressed = Time Interacting with Stressed Conspecific divided by total time interacting: Naive plus Stressed X 100, Fig 2.D). Consistent with the analysis of interaction times, there was a significant Sex by Age by MIA interaction F(1, 150) = 12.05, p = 0.0007, *η*^2^ = 0.05) supporting the conclusion that Saline offspring exhibited preference for PN30 juveniles and avoided PN50 adults, but male Poly I:C offspring did not have preference. Thus, MIA appeared to selectively interfere with social affective preferences in male, but not female, rats.

**Figure 2:**
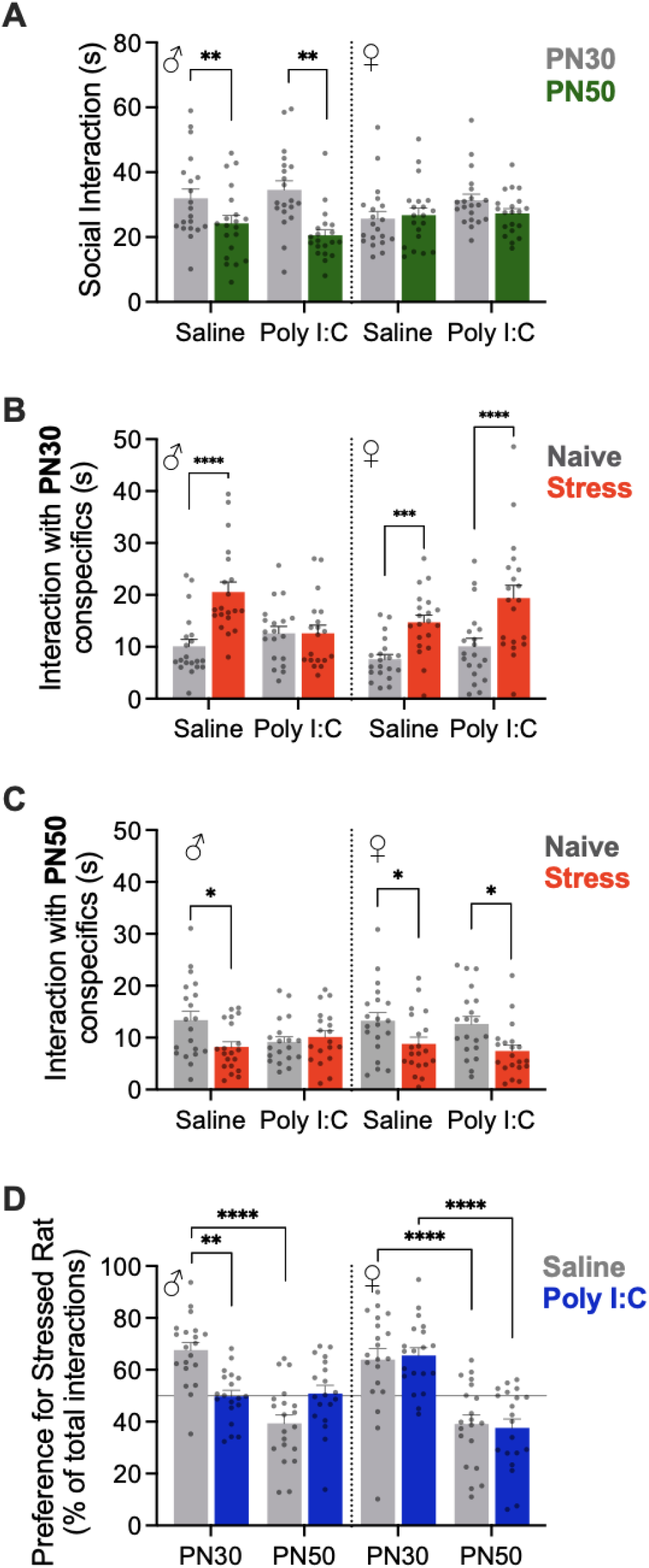
Maternal immune activation altered social affective behavior. **A**. Mean (+/-SEM) time spent interacting with naive conspecifics on day 2 of the SAP test. Males spent more time investigating naive juveniles (PN30) compared to naive adults (PN50), \*\**p*s < 0.01 Sex by Age interaction. **B-C**. Mean (+/-SEM) time spent interacting with naive and stressed juvenile conspecifics on Day 3 of the SAP test with juvenile (**B**) or adult (**C**) conspecifics. Male and Female control rats (saline) and Female MIA rats (Poly I:C) spent more time investigating stressed juveniles but Male MIA rats did not differentiate between naive and stressed conspecifics. **D**. Mean (+/-SEM) preference for interaction with the stressed conspecifics. Values greater than 50% indicate more time spent with stressed conspecific; horizontal line at 50% indicates no preference. Poly I:C male offspring did not show preference for PN30 or avoidance of PN50 stressed conspecifics. Dots indicate individual replicates. *p<0.05, **p<0.01, ***p<0.00, ****p<0.0001 Sidak post hoc comparisons; see text for additional statistics.

### Maternal immune activation has sex specific effects on insular cortex CRF sensitivity

CRF increases excitability of the insular cortex and CRF_1_ receptors in the insula are necessary for social behavior in the SAP test in males but not females (Rieger et al., 2022). Therefore, we hypothesized that Poly I:C offspring may have altered sensitivity to CRF in the insula. To test this hypothesis we recorded insular cortex field excitatory postsynaptic potentials (fEPSPs) in acute slices of the posterior insular cortex from adult MIA or control males and females under baseline (aCSF) and bath application of CRF (Fig. 3A-B). In recordings from male slices (Fig. 3C), significant main effects were found for stimulation voltage (F(10, 190) = 915.6, p < 0.0001, *η*^2^ = 0.92), CRF (F(1, 19) = 12.15, p=0.0025, *η*^2^ = 0.008) and a significant CRF by MIA by Stimulation interaction was found (F(10, 190) = 2.757, p = 0.0034, *η*^2^ =0.001). Recordings from male saline offspring exhibited the previously reported effect of CRF increasing insular excitability causing a leftward shift in input/output (I/O) curves (CRF fEPSPs were significantly greater than aCSF at 3.5V (p = 0.049) and 4, 4.5 and 5V (ps < 0.001). However, male MIA offspring showed no change in insular excitability at any stimulation level following application of CRF, indicating a loss of insular sensitivity to CRF after MIA treatment. In recordings from female slices, significant main effects were found for stimulation voltage (F(10, 180) = 746.1, p < 0.0001, *η*^2^ = 0.89), CRF (F(1, 18) = 19.13, p = 0.0004, *η*^2^ = 0.01) and a CRF by Stimulation interaction (F(10, 180) = 15.14, p < 0.0001, *η*^2^ = 0.008). In recordings from female saline offspring (Fig. 3D), CRF did not significantly alter fEPSPs (no aCSF vs CRF comparisons reached significance at any stimulation voltage, ps > 0.17). However, in recordings from female Poly I:C offspring, fEPSPs were greater after CRF at 3 (p = 0.031), 3.5 (p = 0.002), 4, (p < 0.001), 4.5 and 5V (ps < 0.0001). In sum, MIA appeared to render male insula less sensitive to CRF but rendered female insula more sensitive to CRF.

**Figure 3:**
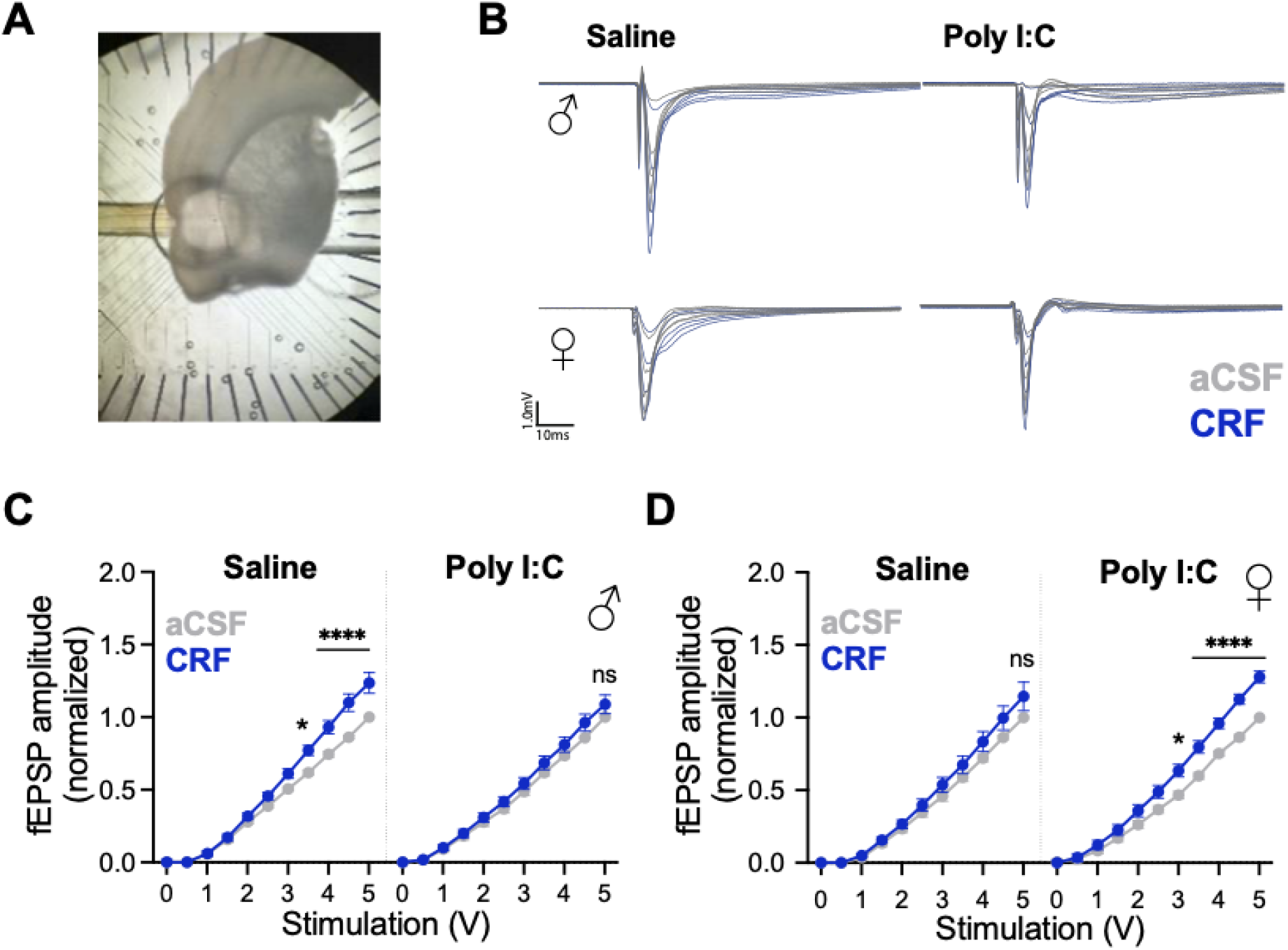
Effect of CRF on insular synaptic efficacy after maternal immune activation in males and females. **A**. Photograph of coronal section with insular cortex placed over the perforated multiple electrode array. **B**. Representative fEPSPs were recorded in aCSF then after CRF (300nM) by biphasic stimuli from 0 to 5V. Data were analyzed as peak fEPSP amplitude normalized to the 5V response at each channel observed in aCSF. **C-D**. Mean (+/-SEM) fEPSP amplitude before and after CRF application in acute slices from adult males (**C**) and females (**D**). In slices from male saline offspring and from female Poly I:C offspring CRF increased fEPSP amplitude at higher voltages (* indicate significant differences between aCSF and CRF fEPSP, see text for additional statistics). CRF had no effect on slices from Poly I:C male or saline female offspring.

### Insular cortex CRF infusion increases social exploration in MIA females but not males

To determine if the gain or loss of insular sensitivity to CRF observed in fEPSPs was behaviorally relevant, we tested whether microinjections of CRF (0.5μL/side at 300nM) to the insula led to changes in one-on-one social exploration of naive (unstressed) juvenile isosexual conspecifics in the offspring of saline or Poly I:C treated dams (Fig. 4A). Similar to our previous result (Rieger et al., 2022), in control males, insular CRF injection led to an increase in social exploration of a naive juvenile but CRF had no effect on control females (Fig. 4B) which was reflected in a significant Sex by MIA by CRF interaction (F(1, 40) = 30.71, p < 0.0001, *η*^2^ = 0.05).

**Figure 4:**
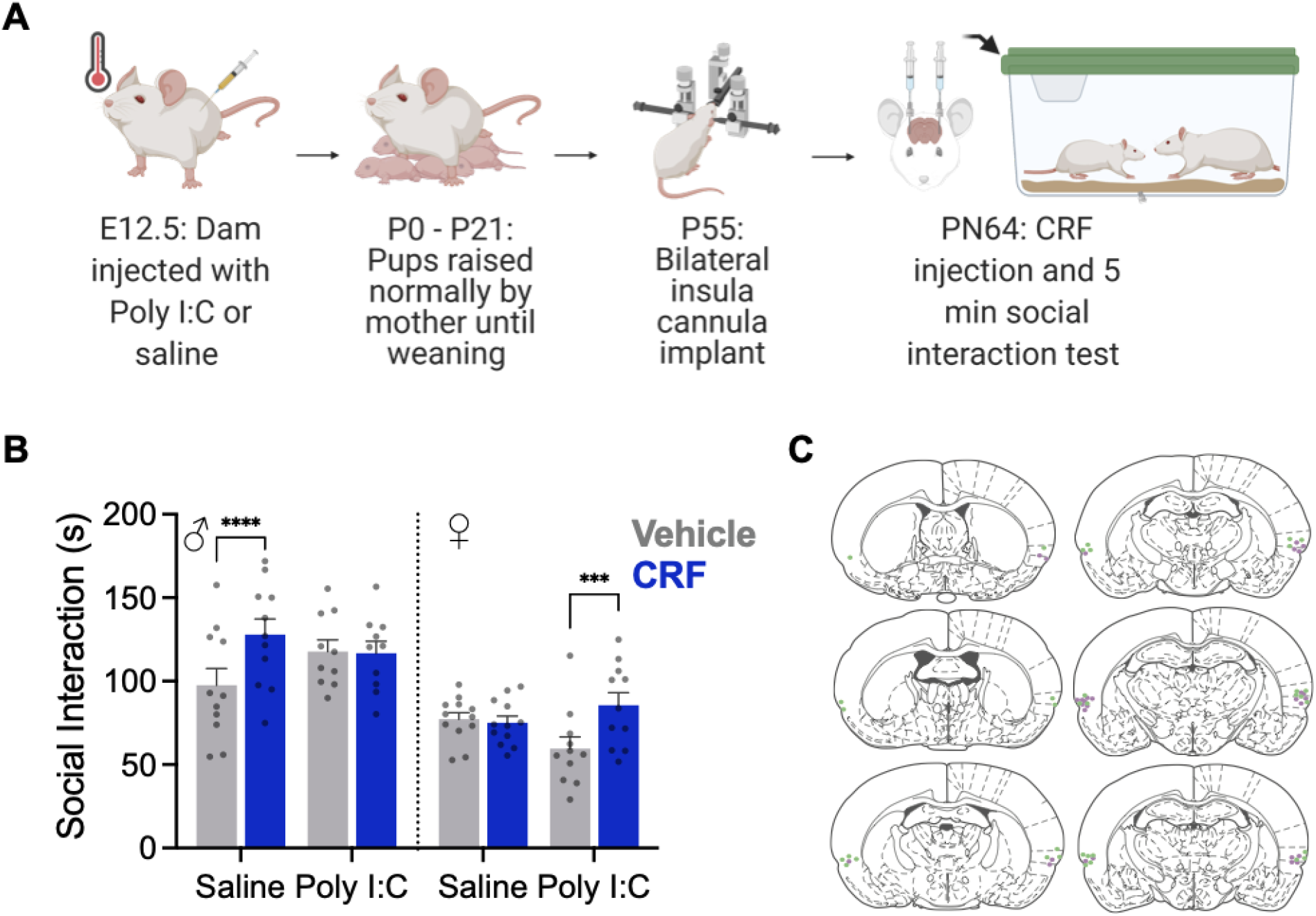
Effect of CRF on social interaction after maternal immune activation in males and females. **A**. Diagram of experimental procedure. **B**. Mean (+/-SEM with individual replicates) time spent interacting with a naive juvenile conspecific in a 5 min social interaction test. Male offspring of saline treated dams and female offspring of Poly I:C treated dams spent more time investigating the juvenile after CRF injection to the insular cortex (***p<0.001, ****p<0.0001). **C**. Representative locations of insula cannula implants.

There was also a significant main effect of Sex with females engaging in less social interaction across conditions (F1, 40) = 30.34, p < 0.0001, *η*^2^ = 0.40). As in the fEPSP result, MIA males showed no change in social exploration after CRF injection whereas CRF increased social interaction in MIA females (Fig. 4B). To summarize, CRF influenced social interaction in control males, not control females. If males were the offspring of a MIA mother, social interaction was no longer influenced by CRF and sensitivity to CRF was decreased. On the other hand, MIA females displayed CRF influence over social interaction compared to their control counterparts, and increased sensitivity to CRF.

### CRF1 receptor distribution is not altered in insular cortex

A functional difference in sensitivity to CRF following MIA could be the consequence of changes at many levels of CRF signaling including, but not limited to, the location, cellular distribution or density of CRF receptors; tonic or phasic characteristics of CRF release; or in any aspect of the G-protein mediated cellular signaling cascades initiated by CRF receptor binding. Indeed, there are many examples of region and sex-specific differences in the CRF system (Bangasser and Wiersielis, 2018). To begin to understand how the CRF system may be affected by MIA, we used fluorescent *in situ* hybridization to determine the distribution of CRF_1_ (*crhr1*), glutamate (*vGlut1*) and GABA (*gad1*) mRNA in the insula (Fig. 5A). Our analysis also included two regions found in the same coronal plane as the posterior insula: the central amygdala, which is implicated in CRF modulation of many behaviors, and the primary somatosensory cortex, which was included as a control region not directly involved in socioemotional behaviors. In each region, we counted the number of mRNA positive cells (those colocalized with DAPI) and the number of putative glutamate and GABA neurons that colocalized with CRF_1_. CRF_1_ mRNA was prevalent in all regions of interest (Fig. 5B) with ∼25% of cells in the insula and CeA positive for CRF_1_ and 35-50% of cells in the sensory cortex. In sections from Poly I:C treated offspring there were more CRF_1_ cells in the sensory cortex (Main effect of MIA, F(1, 28) = 7.027, P=0.013, *η*^2^ = 0.018). This difference appeared to be carried in the males in which Poly I:C counts were significantly greater than saline counts (p = 0.022). The distribution of vGlut1 cells (Fig. 5C) varied by region with 35-40% of cells in the insula, ∼0% of cells in CeA and 35-45% of cells in the sensory cortex. Interestingly, in sections from Poly I:C treated offspring, there was a higher number of vGlut1 positive cells (Main effect of MIA: F(1, 28) = 6.758, p = 0.017, *η*^2^ = 0.018). The distribution of gad1 cells (Fig. 5D) also varied by region with the insula and sensory cortex containing ∼10-15% gad1 cells and the CeA containing 35-45%. Gad1 counts were higher in the sensory cortex of Poly I:C treated sections (Main effect of MIA, F(1, 28) 6.402, p = 0.017, *η*^2^ = 0.018).

**Figure 5:**
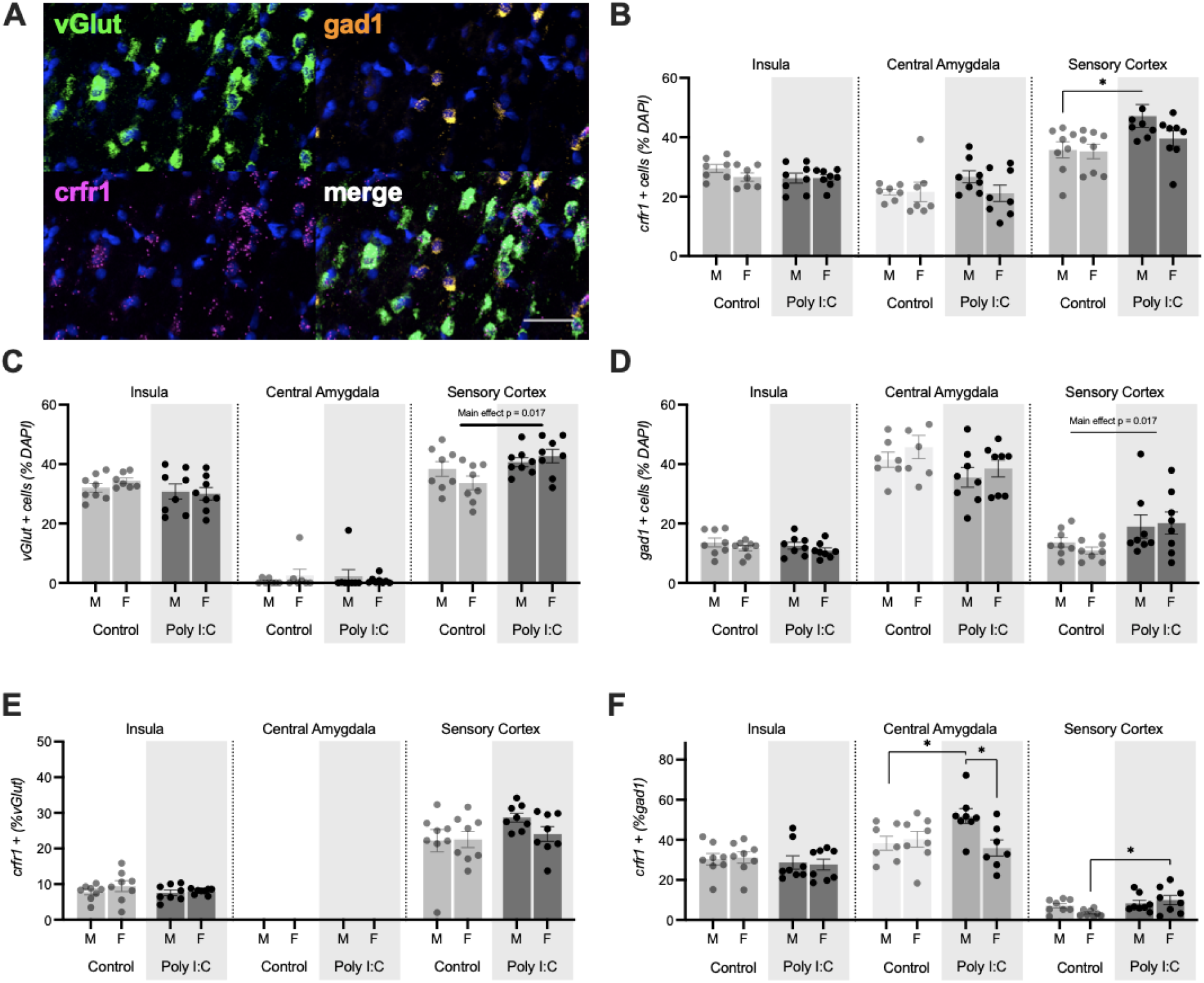
Distribution of CRF_1_ mRNA in insula, central amygdala and sensory cortex. **(A)**. CRF_1_, vGlut1, and gad1 mRNAs were visualized with fluorescent multiplex *in situ* hybridization in male and female adult offspring of control or MIA treated mothers, n = 8 treatment. Composite image from posterior insular cortex. Scale bar = 50μm. **(B)** Percentage of cells in the regions of interest expressing CRF_1_. MIA increased the number of CRF_1_ cells in the sensory cortex (p = 0.013). **(C)**. Percentage of cells in the region of interest expressing vGlut1. MIA increased the number of vGlut1 cells in the sensory cortex (p = 0.017). **(D)** Percentage of cells in the region of interest expressing gad1. MIA increased the number in the sensory cortex (p = 0.017). **(E)** Percentage of vGlut1 neurons coexpressing CRF_1_. **(F)** Percentage of gad1 neurons coexpressing CRF_1_. MIA increased the number of CRF_1_+gad1 neurons in the CEA of males. MIA increased the number of CRF_1_+gad1 neurons in the sensory cortex of females. Bars depict the mean +/-SEM with individual replicates. *p<0.05.

Turning to the analysis of colocalization, CRF1 was found colocalized with vGlut1 (Fig. 5E) and gad1 (Fig. 5F) in most regions of interest. As expected, the glutamatergic and GABAergic markers vGlut1 and gad1 did not colocalize (data not shown). In the insula, about 5-10% of vGlut positive cells contained CRF_1_ and in the sensory cortex ∼25% of cells contained CRF_1_; there were no significant effects of Sex or MIA in either region. CRF_1_ was found with ∼30% of insula gad1 neurons, 40-50% of CeA gad1 neurons and < 10% of sensory cortex gad1 neurons. In the CeA, there were significantly more CRF_1_ positive gad1 neurons in sections from Poly I:C treated male offspring (Sex by MIA interaction, F(1, 26) = 5.593, p = 0.026, *η*^2^ = 0.15), which differed from both control male and female Poly I:C counts (ps < 0.05). In the sensory cortex there was a significant effect of MIA increasing the number of gad1 cells with CRF_1_ (Main effect of MIA, F(1, 28) = 6.326, p = 0.018, *η*^2^ = 0.17); this effect was most prevalent in females which differed across control and Poly I:C conditions (p = 0.026). To summarize, we did not observe any significant effects of Poly I:C treatment or sex on mRNA distribution or colocalization in the insular cortex. MIA increased the number of vGlut, gad1 and CRF_1_ cells in the sensory cortex but reduced the number of gad1 cells in the central amygdala. Sex specific effects of MIA were present in in the CeA where Poly I:C increased the number of CRF_1_ + gad1 cells in males and in the sensory cortex where there was an increase in CRF_1_ + gad1 cells in females.

## Discussion

In this study, we tested for alterations in social decision making by male and female rats exposed to MIA. Using the social affective preference test, we predicted that MIA exposed offspring would both fail to form preferences for novel juvenile and adult conspecifics based on stress cues and show decreased overall sociability. Finally, we hypothesized that this effect would be dependent on CRF sensitivity and CRF_1_ receptor distribution in the insular cortex. We found that MIA male offspring, but not female offspring, failed to develop a social affective preference for both adult and juvenile conspecifics. MIA males also showed a loss of insular sensitivity to CRF, shown first by a loss of response to CRF in insular cortex slices and second by a loss of increased social exploration following CRF injections in the insula. However, MIA female offspring showed an increased sensitivity to CRF compared to controls in both slice physiology and behavioral experiments. Taken together these findings indicate a sex specific change in social decision making and CRF sensitivity following MIA.

Although MIA is reported to reduce sociability itself, general sociability, as determined by total time interacting with conspecifics during the SAP test, was not affected by MIA. This could be a consequence of the dose and timing of Poly I:C administration which are critical considerations in the design and interpretation of MIA studies (Kentner et al., 2019). The dose used here was selected based on a pilot study designed to determine a dose that produced a robust acute inflammatory response (indicated by fever, Fig. 1) without affecting dam survival, litter size and later maternal grooming. In our hands, 0.5mg/kg met this requirement but it falls on the lower end of the range of doses used in other groups. For example, 4mg/kg was used leading to large effects on numerous measures in rats, including females (Osborne et al., 2019) and even higher doses are used in mice (e.g., 20mg/kg in Zhao et al., 2021). Therefore, the MIA treatment used here may model the effect of a relatively mild maternal infection in which later effects are only evident in the approach/avoidance decision present in the social affective preference test. Furthermore, with increasing doses, MIA would likely impair social choice in female offspring. While future studies may consider using higher doses, the current results highlight the opportunity for discovery afforded by expanding the range of behavioral assays that may translate to disease pathophysiology.

We targeted the CRF system in this study because CRF is important for both social decision making (Rieger et al., 2022) and the social transfer of stress (Sterley and Bains, 2021; Sterley et al., 2018) -- two processes which we believe are prime targets to be affected by MIA which alters sociability and stress reactivity (Carlezon et al., 2019; Howerton and Bale, 2012; Zhao et al., 2021). Little is known about how the CRF system and CRF receptor distribution is affected by MIA (Perry et al., 2021). Thus, we studied the insular cortex, an important node for integrating external and internal stimuli (Gogolla, 2017; Rogers-Carter et al., 2018) that is broadly connected to the social decision making network (Gehrlach et al., 2020; Rogers-Carter and Christianson, 2019) and a novel target for MIA studies. Here we find that, in males who were exposed to MIA, the insula, previously shown to be necessary for the formation of social affective preference (Rieger et al., 2022), is less sensitive to CRF causing a loss of social affective preference. This adds to a body of literature on MIA models that indicates that males are more susceptible to prenatal stressors than females (Carlezon et al., 2019; Howerton and Bale, 2012). However, we also show for the first time that not only are females buffered from the overall behavioral deficits of MIA, but also that females show increased sensitivity to CRF in the insula, providing a potential mechanism for how this social deficit buffering occurs. As a loss of insular activation in males due to a loss of sensitivity to neurotransmitters such as CRF causes behavioral deficits, finding ways to activate the insula to rescue the ability to complete social decision making may provide a new avenue for therapeutic research.

Interestingly, despite a change in CRF sensitivity in the insula, no apparent changes in CRF_1_ receptor cells, location or distribution were detected in the insular cortex. We focused on CRF_1_ because it is the predominant CRF receptor in the insular cortex (Li et al., 2002; Potter et al., 1994; Sanchez et al., 1999) and thus the most likely to show an effect in this approach. Because no changes were evident at the level of CRF_1_ mRNA distribution in the insular cortex, it is likely that the effects we see here are tied to something more complex than simply the total amount of CRF_1_. One possibility is that, while MIA is not altering CRF_1_ mRNA it could be leading to desensitization or internalization of CRF_1_ receptors which would not be captured by RNAscope. In the presence of a large amount of agonist, CRF_1_ receptors can become desensitized through G-protein coupled receptor kinase or β-arrestin mechanisms (Hauger et al., 2009). As such, if males exposed to MIA produce high levels of CRF or they maintain higher levels of CRF after stress (Zhao et al., 2021) this could lead to an overall desensitization of the CRF receptor preventing signaling. CRF receptors can also become internalized as a result of high agonist levels (Reyes et al., 2006), which would prevent CRF signaling but not necessarily be captured by mRNA levels. While we found no changes in CRF_1_ receptor distribution in the insula we did find that CRF_1_ was increased in the sensory cortex of males and CRF_1_ had greater colocalization with GABA cells in the central amygdala. Importantly the insula has strong bidirectional connectivity with the sensory cortex and strong outputs to the central amygdala (Gehrlach et al., 2020). As such, the effects we observed may be indicative of broader alterations in the way CRF modulates stress and affective processes in brain regions affected by MIA. On alternative builds on the discovery that MIA alters the regulation of corticosterone in social encounters, with sustained corticosterone observed compared to controls (Zhao et al., 2021). Therefore, altered CRF production, drive or impaired negative feedback of the hypothalamic stress response are intriguing complementary processes which may contribute to reduced CRF_1_ sensitivity in insula and loss of social affective preference.

It is also possible that the CRF system alone is not driving the behavioral changes we see in the MIA exposed males of this study. Our previous work in non MIA rats has shown that endocannabinoids are necessary for CRF activation of the insular cortex via retrograde action at CB_1_ receptors causing presynaptic inhibition (Rieger et al., 2022) and other work has shown that eCB signaling is dysregulated following MIA (Osborne et al., 2019). Because CRF_1_ receptor distribution does not change it could instead be that eCB production or CB_1_ receptor distribution does change in the insula following MIA. In this case CRF sensitivity would be decreased due to CB_1_ receptors not being activated and presynaptic inhibition remaining in place, stopping the insula from activating in response to conspecifics. In this study we did not test for changes in CB_1_ in MIA animals and therefore it could be that changes in CB_1_ distribution account for the alterations seen in CRF signaling.

Overall we find that, similar to previous studies (Bale et al., 2010; Bilbo and Schwarz, 2012; Carlezon et al., 2019; Haida et al., 2019; Zhao et al., 2021), MIA leads to sex specific changes in behavior. However, in this study, we also show that social decision making - in the form of assessing stress cues and the ability to form preferences towards novel conspecifics - is lost in males but not females exposed to MIA, while sociability is maintained in both sexes. We also show, for the first time, that both the social deficit seen in males and the buffering seen in females may be due to changes in CRF signaling sensitivity with males losing but females gaining sensitivity after exposure to MIA. Insular cortex is a hub of affective and salience processes (Gogolla, 2017, Shin & Liberzon, 2010; Paulus & Stein, 2006; Uddin & Mennon, 2009) that are disrupted in a wide range of psychopathologies from autism spectrum disorder, substance use disorders, eating disorders, depression, schizophrenia and anxiety. Early life and maternal stressors all increase the risk for developing these conditions so these new findings take a step towards a more complete understanding of how maternal sickness shapes the offspring behavior and new avenues of translational study.

## Author Contributions

Conceptualization, N.S.R., J.P.C.; Methodology, N.S.R., J.P.C.; Investigation, N.S.R., A.J.N., S.L., B.H.B., J.P.C.; Writing – Original Draft, N.S.R. and J.P.C.; Writing -- Revision & Editing, N.S.R. and J.P.C.; Funding Acquisition, J.P.C. and NSR.

## Acknowledgement

The authors wish to thank Dr. Bret Judson, director of the Boston College Imaging Core, for training and assistance with all microscopy, Nancy McGilloway and Todd Gaines, administrators of the Boston College Animal Care Facility, for outstanding animal husbandry.

